# Inhibitory effect of lactobacilli supernatants on biofilm and filamentation of *C. albicans, C. tropicalis*, and *C. parapsilosis*

**DOI:** 10.1101/2022.11.26.518025

**Authors:** Yeuklan Poon, Mamie Hui

## Abstract

Probiotic *Lactobacillus* strains had been investigated for the potential to protect against infection caused by the major fungal pathogen of human, *Candida albicans*. Besides antifungal activity, lactobacilli demonstrated a promising inhibitory effect on biofilm formation and filamentation of *C. albicans*. On the other hand, two commonly isolated non-albicans *Candida* species, *C. tropicalis* and *C. parapsilosis*, have similar characteristics in filamentation and biofilm formation with *C. albicans*. However, there is scant information of the effect of lactobacilli on the two species. In this study, biofilm inhibitory effects of *L. rhamnosus* ATCC 53103, *L. plantarum* ATCC 8014, and *L. acidophilus* ATCC 4356 were tested on the reference strain *C. albicans* SC5314 and six bloodstream isolated clinical strains, two each of *C. albicans, C. tropicalis*, and *C. parapsilosis*. Cell-free culture supernatants (CFSs) of *L. rhamnosus* and *L. plantarum* significantly inhibited *in vitro* biofilm growth of *C. albicans* and *C. tropicalis. L. acidophilus*, conversely, had little effect on *C. albicans* and *C. tropicalis* but was more effective on inhibiting *C. parapsilosis* biofilms. Neutralized *L. rhamnosus* CFS at pH 7 retained the inhibitory effect, suggesting that exometabolites other than lactic acid produced by the *Lactobacillus* strain might be accounted for the effect. Furthermore, we evaluated the inhibitory effects of *L. rhamnosus* and *L. plantarum* CFSs on the filamentation of *C. albicans* and *C. tropicalis* strains. Significantly less *Candida* filaments were observed after co-incubating with CFSs under hyphae-inducing conditions. Expressions of six biofilm-related genes (*ALS1, ALS3, BCR1, EFG1, TEC1*, and *UME6* in *C. albicans* and corresponding orthologs in *C. tropicalis*) in biofilms co-incubated with CFSs were analyzed using quantitative real-time PCR. When compared to untreated control, the expressions of *ALS1, ALS3, EFG1*, and *TEC1* genes were downregulated in *C. albicans* biofilm. In *C. tropicalis* biofilms, *ALS3* and *UME6* were downregulated while *TEC1* was upregulated. Taken together, the *L. rhamnosus* and *L. plantarum* strains demonstrated an inhibitory effect, which is likely mediated by the metabolites secreted into culture medium, on filamentation and biofilm formation of *C. albicans* and *C. tropicalis*. Our finding suggested an alternative to antifungals for controlling *Candida* biofilm.

## 1 Introduction

*Candida* species are commensal fungi that colonize healthy human skin, gastrointestinal and genital tracts. They are also opportunistic pathogens, leading to mucosal and systemic infections(Kim and Sudbery, 2011). *Candida* are one of the most important agents of fungal infection worldwide and the most frequently isolated fungal species in healthcare-associated infections(Brown et al., 2012). Over 90% of invasive *Candida* infections were caused by five species, *Candida albicans, C. glabrata, C. tropicalis, C. parapsilosis*, and *C. krusei*, with *C. albicans* accounted for 47-66% of cases (Pfaller and Diekema, 2007; Pfaller et al., 2019). Being the predominant cause of fungal infection, *C. albicans* has become the best-studied *Candida* species(Kabir et al., 2012). Although *C. albicans* remains as the most common isolated single species, a progressive shift to increased prevalence of non-albicans *Candida* species has been observed over the past decades (Pfaller et al., 2019). In the global SENTRY antifungal surveillance program, the frequency of *C. albicans* was reported to gradually decrease from 57.4% of invasive *Candida* isolates in 1997-2001 to 46.4% in 2015-2016(Pfaller et al., 2019).

Compared to *C. albicans*, the emerging non-albicans species exhibit reduced antifungal susceptibility. For example, *C. krusei* is considered as intrinsically resistant to fluconazole. *C. glabrata* has reduced susceptibility to azoles, while *C. parapsilosis* shows elevated minimum inhibitory concentration to echinocandins(CLSI, 2008). Among the isolates collected in 2006-2016 for the SENTRY Program, the fluconazole resistance rates of *C. albicans* was 0.3%, while that of *C. glabrata, C. parapsilosis*, and *C. tropicalis* were 8.1%, 3.9%, and 3.2% respectively(Pfaller et al., 2019). For echinocandins, the resistance rates were low in *C. albicans* (0-0.1%) and *C. parapsilosis* (0-0.1%) but were higher in *C. tropicalis* (0.5-0.7%), *C. krusei* (0-1.7%), and *C. glabrata* (1.7-3.5%).

Among the four common isolated non-albicans species, *C. tropicalis* and *C. parapsilosis* are phylogenetically closely related to *C. albicans*(Butler et al., 2009). Similar to *C. albicans, C. tropicalis* and *C. parapsilosis* can undergo yeast-filament transition, which is an important virulence factor of *C. albicans* for invading host epithelial cells and causing tissue damage(Sudbery, 2011). *C. albicans* and *C. tropicalis* produce true hyphae and pseudohyphae while *C. parapsilosis* produces only pseudohyphae(Lackey et al., 2013).These three species are also capable of forming biofilm composed of yeast cells and filaments.

*Candida* biofilm formed *in vivo* on abiotic surfaces of indwelling medical devices, such as dentures, endotracheal tubes, prosthetic device, and various types of catheters, can lead to device failure and infections, including fatal catheter-related bloodstream infections(Ramage et al., 2006). Moreover, *Candida* biofilm has shown significantly decreased susceptibility to some commonly used antifungal agents, such as fluconazole and amphotericin B, when compared to the planktonic cells(Ramage et al., 2001; Pierce et al., 2008). Biofilm protects *Candida* cells from antifungal agents, leading to persistent infection that is difficult to eliminate without removal of the infected implant(Ramage et al., 2006).

In view of the emergence of antifungal resistance, the use of probiotic *Lactobacillus* species has been suggested as an alternative therapeutic option against *Candida* infection(Matsubara et al., 2016a). Several clinical trials reported that the administration of *Lactobacillus* improve outcome of vaginal candidiasis and reduce colonization of *Candida* in oral cavity of elderly people and in gastrointestinal tract of preterm neonates(Roy et al., 2014; Kovachev and Vatcheva-Dobrevska, 2015; Kraft-Bodi et al., 2015). *In vitro* studies also demonstrated that lactobacilli possess anticandidal ability, inhibitory effect on adhesion to host cells, and inhibitory effect on biofilm formation and filamentation of *C. albicans*(Strus et al., 2005; Köhler et al., 2012; Coman et al., 2015; Vilela et al., 2015; Matsubara et al., 2016b; Song and Lee, 2017). Interestingly, as demonstrated with cell-free supernatants and in agar overlay assays, direct interaction of lactobacilli with *C. albicans* cells is unnecessary for the inhibitory effect, suggesting that the effect is exhibited by the metabolites produced by lactobacilli(Matsubara et al., 2016b; Liao et al., 2019).

Although the effect of lactobacilli on *C. albicans* has been extensively studied, there is a relative lack of studies on non-albicans species. The aim of this study was to investigate the inhibitory effect of *L. rhamnosus, L. plantarum*, and *L. acidophilus* on the biofilm formation of *C. albicans* and closely related non-albicans species, *C. tropicalis* and *C. parapsilosis*. In addition, we evaluated the change in filamentation and the expression of filamentation- and biofilm-related genes of *C. albicans* and *C. tropicalis* when incubated with cell-free supernatants of *L. rhamnosus* and *L. plantarum*.

## 2 Materials and Methods

### 2.1 Strains and Culture Conditions

Six *Candida* clinical strains (*C. albicans* A14 and A69, *C. tropicalis* T18S and T38R, and *C. parapsilosis* P149 and P152) originally isolated from bloodstream specimens, and a laboratory reference strain *C. albicans* SC5314 were used. All strains were identified with MALDI-TOF mass spectrometry with MALDI Biotyper (Bruker, USA). Prior to experiment, *Candida* cells were first subcultured from glycerol stock culture onto Sabouraud dextrose agar (SDA) and incubated at 35 °C for at least 1 day to assess the purity and viability of the culture.

Type strains *L. rhamnosus* GG ATCC 53103 (LGG), *L. plantarum* ATCC 8014 (LP8014), and *L. acidophilus* ATCC 4356 (LA4356) were used. Stock cultures were prepared by resuspending bacterial cells in de Man, Rogosa and Sharpe (MRS) broth with 10% glycerol and stored at -80 °C until use. Before experiments, the bacterial cells were first streaked from stock culture onto MRS agar and incubated at 37°C under microaerophilic conditions (5% O_2_, provided by CampyGen, Thermo Fisher Scientific, USA) for 2 days.

### 2.2 Biofilm inhibition

*Candida* strains were grown in yeast extract peptone dextrose (YPD) broth at 37 °C with shaking (200 r.p.m.) for 18 h. The cells were harvested by centrifugation, washed with phosphate buffered saline (PBS), and resuspended to 10^6^-10^7^ CFU/ml in RPMI 1640 medium (with L-glutamine, buffered with MOPS; Thermo Fisher Scientific, USA). For *Lactobacillus, Lactobacillus* colonies were collected from MRS plates and inoculated into MRS broth. After incubated microaerophilically at 37 °C with shaking (200 r.p.m.) for 18 h, the liquid cultures were centrifuged to separate spent media and bacterial cells. Cell-free supernatant (CFS) was prepared by filtering the spent media through a membrane filter with a 0.22 μm pore size (Merck Millipore, USA). MColorpHast pH indicator strip (Millipore, USA) was used to measure the pH of CFS. The harvested *Lactobacillus* cells were washed twice with PBS and resuspended in MRS broth to 1 × 10^7^ CFU/ml.

The incubation of biofilm was divided into two parts, adhesion phase and formation phase(Kuhn et al., 2002). Briefly, in adhesion phase, 100 μl of *Candida* inoculum were seeded into the wells of flat bottom 96-well plates. Plates were prepared in duplicate and then incubated statically at 37 °C for 90 mins. After incubation, spent medium was aspirated from the well. The wells were washed with PBS to remove non-adherent cells. In the formation phase, 100 μl of RPMI medium and 50 μl of *Lactobacillus* CFS, living cell suspension, or MRS broth as control were added per wells. There were five replicate wells for each condition per plate. The plates were incubated statically at 37 °C for an additional 24 h. After incubation, a MColorpHast pH strip was used to check the pH of spent medium. The biofilms were quantified by two methods, XTT reduction assay and crystal violet staining.

For XTT reduction assay, 100 μl of 2,3-Bis-(2-Methoxy-4-Nitro-5-Sulfophenyl)-2H-Tetrazolium-5-Carboxanilide (XTT, 0.5 mg/ml in PBS, Sigma-Aldrich, USA) with 1 μM of menadione (Sigma-Aldrich, USA) were added per well(Pierce et al., 2008). The plate was incubated in dark at 37°C for 2.5h. The resulting-colored supernatant (80 μl) was transferred to a round bottom 96-well plate and its absorbance was measured at 492 nm using a LED microplate reader (Ledetect 96, Labexim Products, Austria). For crystal violet staining, the biofilm in the well was first fixed with 100 μl of 99% methanol, then stained with 100 μl of 0.0625% w/v crystal violet (CV) solution(Peeters et al., 2008). To solubilize the stain, 150 μl of 33% acetic acid were added per well. The solution (100 μl) was transferred to a round bottom plate and its absorbance was measured at 570 nm(Silva et al., 2009).

To examine whether a lower pH contributed to the inhibitory effect of CFS, the assay was repeated using MRS broth acidified to pH 4.0 with lactic acid, and LGG CFS neutralized to pH 7.0 with sodium hydroxide.

### 2.3 Filamentation inhibition

The *C. albicans* and *C. tropicalis* strains were tested against the CFSs of lactobacilli LGG or LP8014, which were prepared from 18h-old culture as previously described and stored at -20°C until use. MRS broth was used as control. The assay was performed following the method described by Wang *et al*. with some modifications(Wang et al., 2017). In a 1.5 ml Eppendorf tube, 144 μl of *Candida* suspension in RPMI medium at concentration of 1 × 10^7^ CFU/ml were mixed with 300 μl of *Lactobacillus* CFS or MRS broth and then topped up to a total volume of 900 μl with RPMI. Additional 10% v/v of fetal bovine serum were added for *C. tropicalis* strains to induce filamentation. After incubating statically at 37 °C for 4 h, the mixture was vortexed for 20 s and 20 μl of it were loaded into a Fuchs-Rosenthal counting chamber (Marienfeld, Germany). Microphotographs of ten 0.0625 mm^2^ squares of the chamber were captured using a AmScope MD500 digital eyepiece camera (AmScope, USA) under a light microscope with a 10× objective lens. The numbers of filaments within the ten 0.0625 mm^2^ squares were counted from the photographic images.

### 2.4 Gene expression in biofilm

To determine the effects of *Lactobacillus* CFS on the transcription of biofilm-related genes in *C. albicans* and *C. tropicalis*, the gene expression levels of *ACT1, ALS1, ALS3, BCR1, EFG1, TEC1, RIP1*, and *UME6* of *C. albicans* and the corresponding orthologs in *C. tropicalis* were evaluated using quantitative real-time polymerase chain reaction(qPCR). Specific primer pairs were designed using the NCBI/Primer-BLAST tool (https://www.ncbi.nlm.nih.gov/tools/primer-blast/) except *RIP1*, primers for which were adopted from previous literature(Nailis et al., 2006). The sequences of primers were listed in Table S1 and S2.

To extract RNA, biofilms of *C. albicans* and *C. tropicalis* were formed in a flat bottom, cell culture-treated 48-well plate (Thermo Fisher Scientific, USA) using the two-phase procedures described for biofilm inhibition, with the volumes of inoculum and CFS (or MRS broth) increased to 200 μl and 100 μl respectively. After incubated statically at 37 °C for 24 h, RNA was extracted from biofilms using the TRIzol™ Plus RNA Purification Kit (Thermo Fisher Scientific, USA) with homogenization done on a Fastprep-24 5G instrument (MP Biomedicals, USA) with lysing matrix Y 2ml tubes at a speed of 6.0 m/s for 30 s twice, with a cooling interval on ice for 3 minutes. The RNA was eluted with 30 μl of RNase-free water preheated to 60 °C. RNA samples were DNase treated using the TURBO DNA-*free* Kit (Thermo Fisher Scientific, USA) according to the manufacturer’s instruction. Complementary DNA (cDNA) was reverse transcribed from 500 ng of RNA by the High-Capacity cDNA Reverse Transcription Kit with RNase inhibitor (Thermo Fisher Scientific, USA). cDNA samples were stored at -80 °C until use.

qPCR was performed in 10 μl reaction volume using the SsoAdvanced Universal SYBR Green Supermix (Bio-rad, USA) on a StepOnePlus Real-Time PCR System (Thermo Fisher Scientific, USA). *Candida* strain, as well as no template controls, was performed within a single qPCR run (96-well plate) with three technical replicates per condition. The reactions started from a primary denaturation at 95 °C for 30 s, followed by 40 cycles of 95 °C for 15 s and 60 °C for 60 s. Specificity of amplification was confirmed by melting curve analysis. Expression levels of the genes of interest were normalized to the reference genes *ACT1* and *RIP1* using the following formula: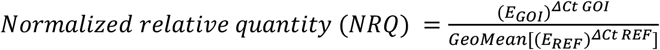, where E represents the amplification efficiency and ΔCt is the difference in cycle threshold (Ct) values of the treatment and control group(Nailis et al., 2006; Hellemans et al., 2007).

### 2.5 Statistical Analysis

Experiments were repeated in triplicate at different occasions. Results of experiment groups in the biofilm inhibition assay were expressed as a percentage relative to the control (MRS) and compared to the control using Student’s t test. Results of filamentation assay were compared to control using one-way analysis of variance (ANOVA) followed by Dunnett’s test. Expression levels of genes in biofilm were expressed as NRQ as described. Student’s t test was used to compare the binary logarithms of the NRQs of experimental group to control group of the same *Candida* strain. A *p*-value ≤ 0.05 was considered significant in all tests. Statistical analyses were performed using Prism (Version 9, GraphPad Software, USA and Microsoft Excel (Office 365, Microsoft, USA).

## 3 Results

### 3.1 Biofilm inhibition

The LGG CFS significantly reduced the biomass of the biofilms produced by five *Candida* strains and the metabolic activity of three strains (Figure 1B). The inhibition was most pronounced (>50% in biomass, P < 0.01) in *C. albicans*, followed by *C. tropicalis*. The LP8014 CFS also shown inhibitory effects to some extent and significantly reduced the biomass and metabolic activity of biofilm formed by *C. albicans* SC5314 (Figure 1D, P < 0.05). However, living LGG and LP8014 cell suspensions had little to no effect (Figure 1A and C). On the other hand, LA4356’s cells significantly reduced both biomass and metabolic activity of the two *C. parapsilosis* strains whiles its CFS also had similar effect (Figure 1E and F).

**FIGURE 1.**
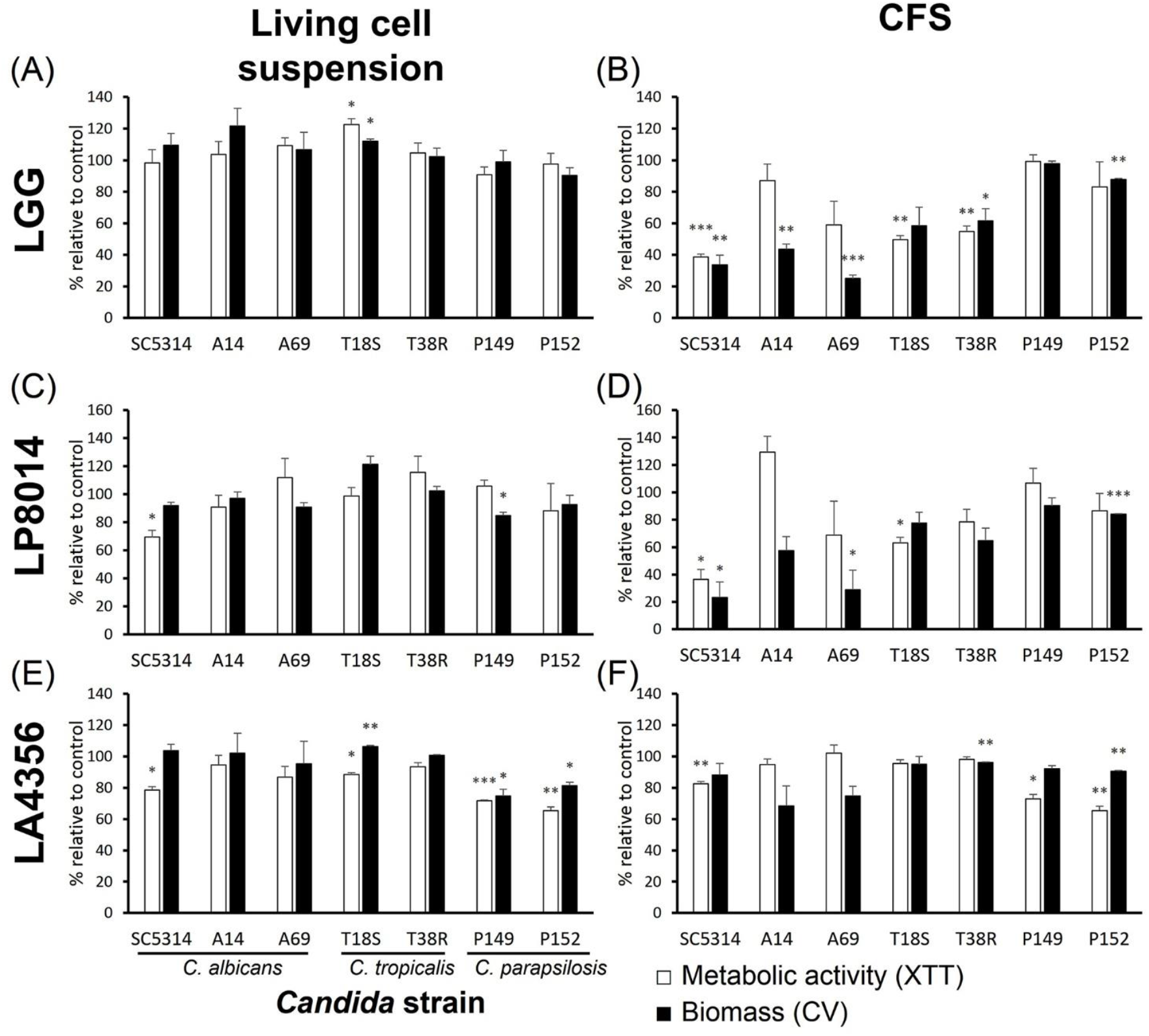
Effects of living cell suspension and CFS of three Lactobacillus strains on biofilm formation of seven *Candida* strains. **(A and B)** *L. rhamnosus* GG, **(C and D)** *L. plantarum* ATCC 8014, **(E and F)** *L. acidophilus* ATCC 4356. Data are shown as percentage relative to paired control (MRS broth) and the mean ± SEM of triplicate experiments. Comparison with control was performed with one-sample t-test. *, p < 0.05; **, p ≤ 0.01; ***, p ≤ 0.001.

Both LGG CFS and LP8014 CFS had a median pH of 4.0, but LA4356 CFS was less acidic (median pH 5). When used in inhibitory assay, the CFSs lowered the median pH of the spent medium after 24-hour incubation from 7.5 (control, MRS broth) to 6.0 (LGG), 6.5 (LP8014), and 7.0 (LA4356) respectively. The medium pH of spent medium was 7.0 in the living cell groups of the three *Lactobacillus* strains. However, when incubated with RPMI medium without *Candida* inoculum, all three *Lactobacillus* strains were able to reduce the medium pH to 4.5 after 24 h.

To examine whether a lower pH contributed to the inhibitory effect of CFS, the assay was repeated with MRS broth acidified to pH 4.0, and LGG CFS neutralized to pH 7.0. As shown in Figure 2, significant inhibition by acidified MRS was only found in biomass of *C. albicans* A14 (35.3% of reduction) and metabolic activity of T18S (12.9%) when compared to MRS control. Neutralized LGG CFS only shown significant reduced inhibitory effect on metabolic activity of *C. albicans* A69 and on biomass of A14 and A69 when compared to untreated LGG CFS.

**FIGURE 2.**
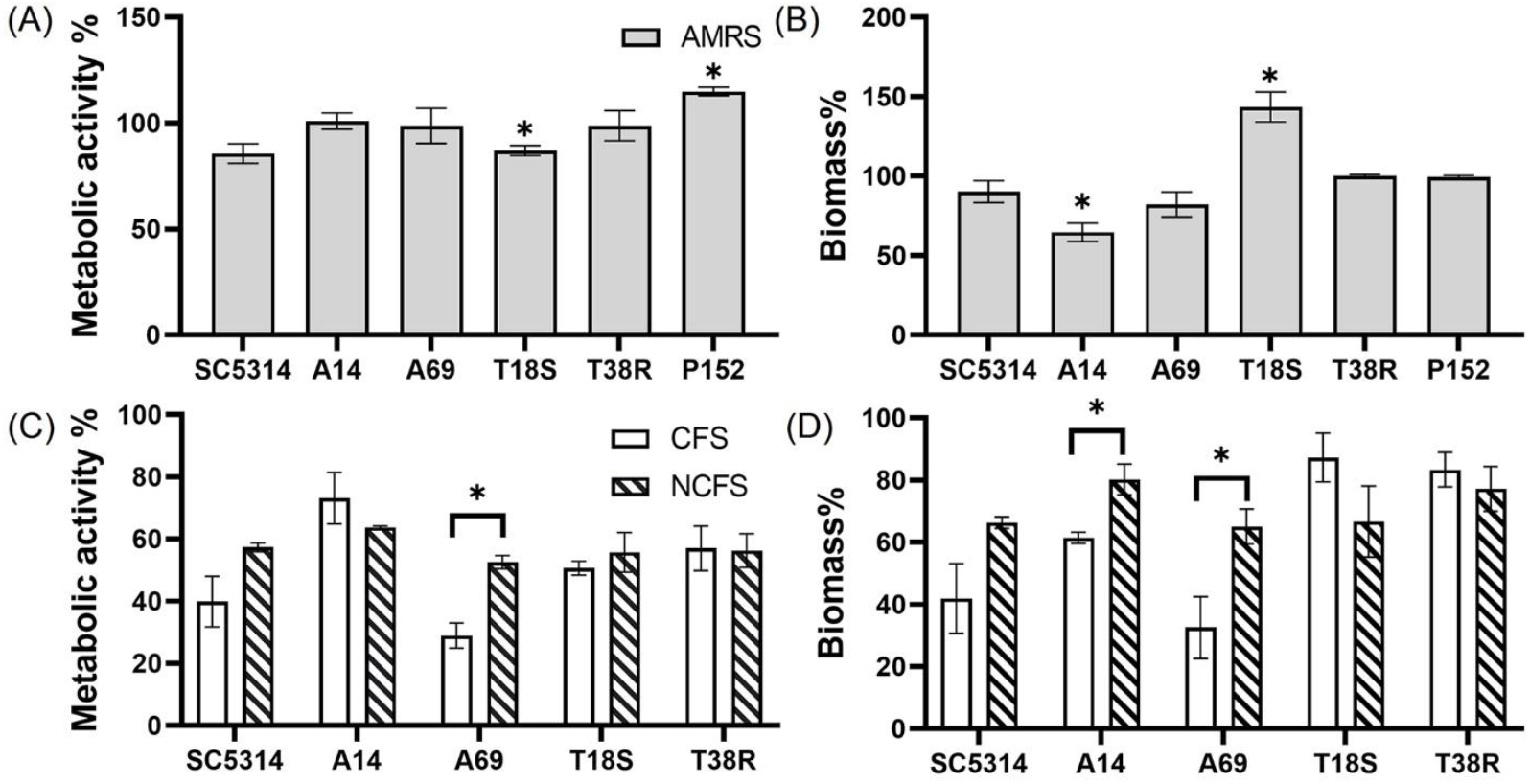
Effects of acidified MRS broth (AMRS) and neutralized CFS (NCFS) on *Candida* biofilm. Data are shown as percentage relative to paired control (untreated MRS) and as mean ± SEM of triplicate experiments.**(A and B)** Effects of AMRS on the metabolic activity and biomass of biofilms produced by three *C. albicans* (SC5314, A14, and A69), two *C. tropicalis* (T18S and T38R), and one *C. parapsilosis* (P152) strains. Asterisks indicate data that are significantly different from control (MRS) at p < 0.05 as determined by one-sample t-test. **(C and D)** Effects of untreated CFS (pH 4.0) and NCFS (pH 7.0) of LGG on the *C. albicans* and *C. tropicalis* strains. Comparison between CFS, NCFS and control (MRS) was done with one-way ANOVA with Tukey’s test. Asterisks indicate significant difference between CFS and NCFS results at p < 0.05.

### 3.2 Filamentation inhibition

As the CFSs of LGG and LP8014 showed pronounced inhibitory effects on biofilms of *C. albicans* and *C. tropicalis*, their effects on the filamentation of *C. albicans* and *C. tropicalis* were further investigated. In the filamentation assay, a large number of *Candida* cells developed into filaments after incubating in RPMI medium mixed with MRS broth (control). However, many *Candida* cells remained in yeast form in the presence of *Lactobacillus* CFSs. Figure 3 showed the examples of cell morphology of each tested *Candida* strains under different conditions. Filamentations of *C. albicans* SC5314, *C. tropicalis* T18S and T38R (Figure 4A, D and E) were statistically significantly inhibited by both LGG CFS and LP8014 CFS (P < 0.05). The number of filaments of *C. albicans* A14 was significantly reduced by LGG CFS (P < 0.05) but not LP8014 CFS when compared to control (Figure 4B). The filamentation of *C. albicans* A69 when incubated with CFSs was inhibited but not significantly different from the control (Figure 4C).

**FIGURE 3.**
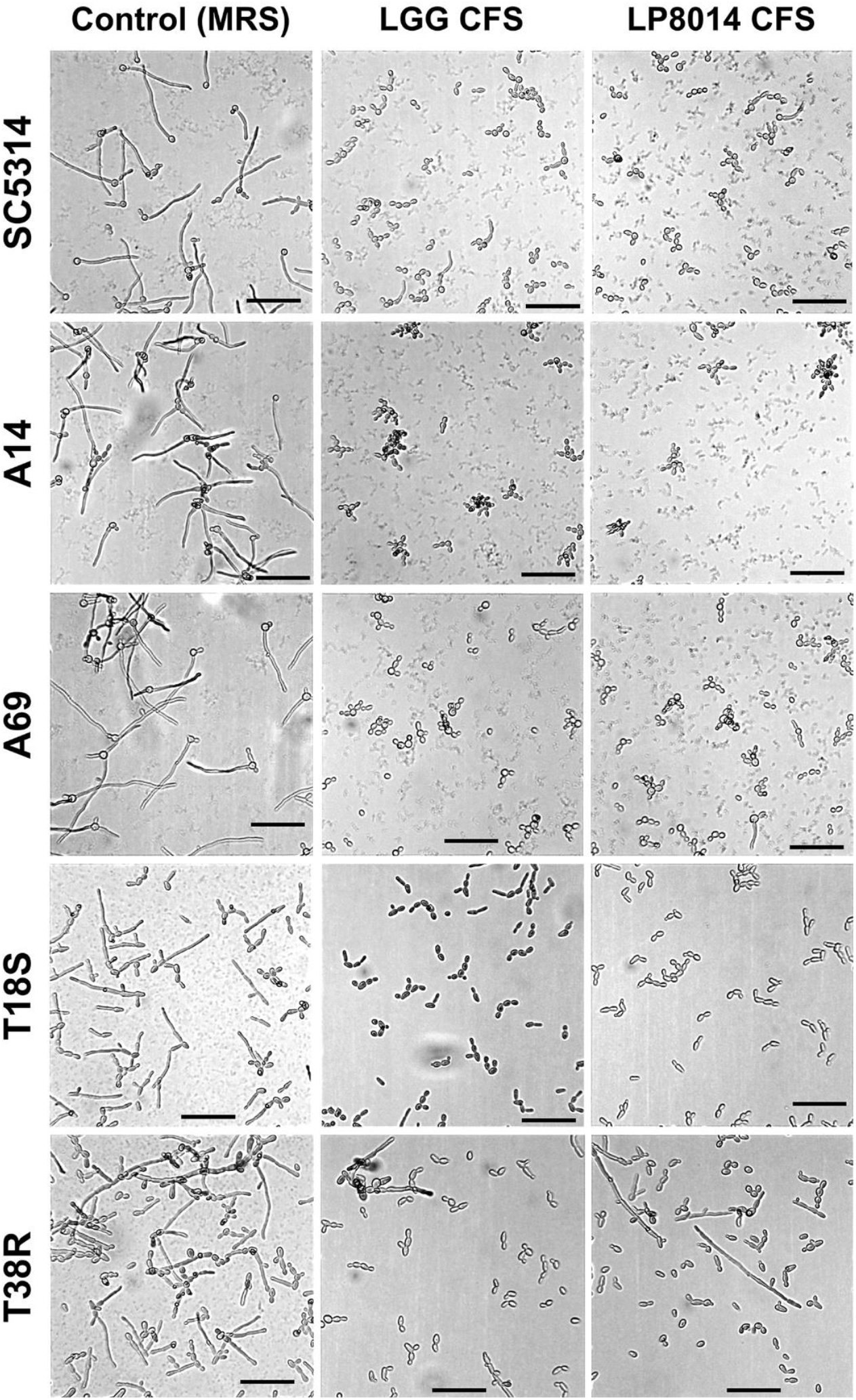
Light micrographs of *C. albicans* and *C. tropicalis* cells in the filamentation assay. In each row, images of one *Candida* strain under three conditions: Control (left), LGG CFS (middle), and LP8014 CFS (right) are shown. Scale-bar: 50 μm.

**FIGURE 4.**
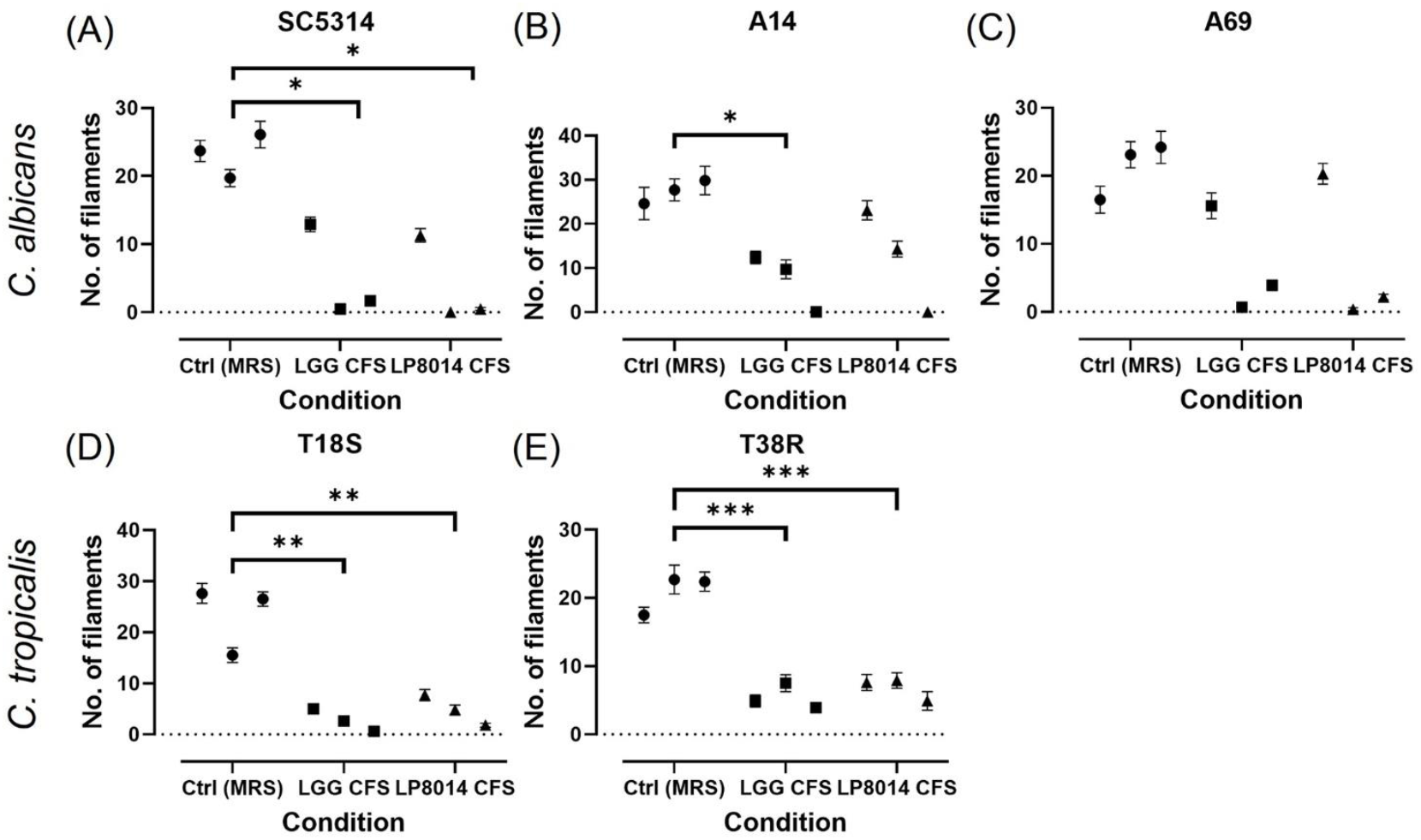
Quantification of filament in the filamentation inhibition assays. **(A)** *C. albicans* SC5314, **(B)** *C. albicans* A14, **(C)** *C. albicans* A69, **(D)** *C. tropicalis* T18S, and **(E)** *C. tropicalis* T38R. Each data point represents the mean ± SEM of the number of filaments in one experiment. Comparisons with the control group were performed with results of triplicate experiments and Dunnett’s test. *, p < 0.05; **, p ≤ 0.01; ***, p ≤ 0.001.

**FIGURE 5.**
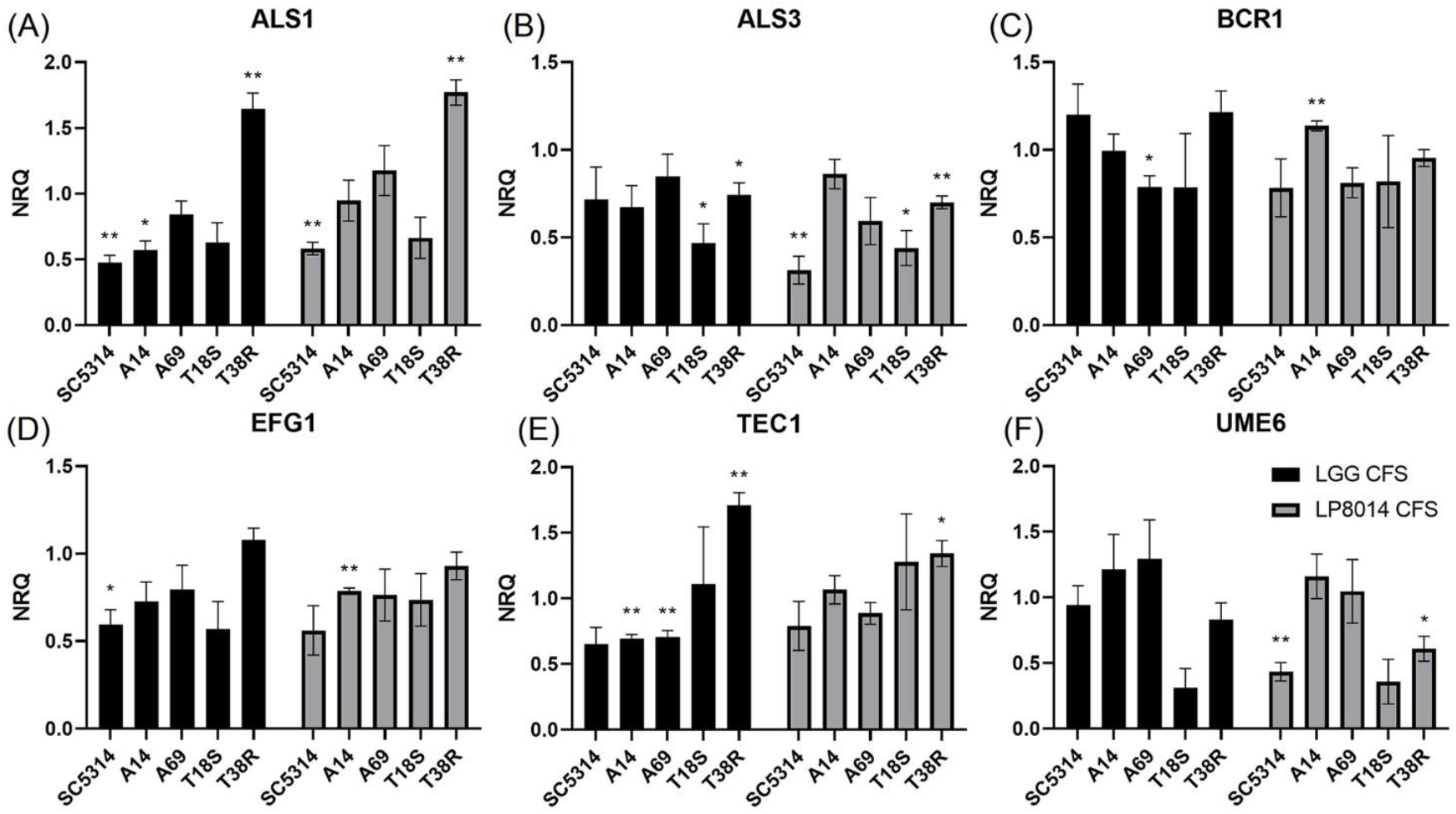
Expression levels filamentation and biofilm-related genes in biofilms of *C. albicans* and *C. tropicalis* incubated with LGG or LP8014 CFSs. (A) *ALS1*, (B) *ALS3*, (C) *BCR1*, (D) *EFG1*, (E) *TEC1*, and (F) *UME6*. Expression levels were expressed as the normalized relative quantity (NRQ). NRQ of the control group (MRS) is constantly 1.0 and not shown on the figures. Values represent the mean ± SEM of triplicate experiments. Comparison with control group was performed with the binary logarithm of NRQ by Student’s t test. *, p < 0.05; **, p ≤ 0.01.

### 3.3 Gene expression in biofilm

Expression of *ALS1, ALS3, BCR1, EFG1, TEC1*, and *UME6* genes in biofilms of *C. albicans* and *C. tropicalis* strains incubated with LGG or LP8014 CFSs were quantified by qPCR. Although being significant only in some of the strains, there were consistent downregulations in *ALS1, EFG1* and *TEC1* genes caused by LGG CFS, and in *ALS3* and *EFG1* genes by LP8014 CFS in the biofilm of the *C. albicans* strains SC5314, A14, and A69 (Figure 8). On the other hand, *ALS3* and *UME6* genes were downregulated and *TEC1* gene was upregulated by both CFSs in *C. tropicalis* T18S and T38R. Interestingly, the expression levels of *ALS1* in T38R were significantly increased to 1.648- and 1.770-fold by both CFSs (Table 1). The expression level of *UME6* gene was downregulated by LP8014 CFS in SC5314, T18S, and T38R and by LGG CFS in T18S, but the differences were not statistically significant.

**TABLE 1.** Expression levels filamentation and biofilm-related genes in biofilms of *C. albicans* and *C. tropicalis* incubated with LGG or LP8014 CFSs.

| Condition | Strain | *ALS1* |  | *ALS3* |  | *BCR1* |  | *EFG1* |  | *TEC1* |  | *UME6* |  |
| --- | --- | --- | --- | --- | --- | --- | --- | --- | --- | --- | --- | --- | --- |
|  |  | NRQ* | *P*† | NRQ | *P* | NRQ | *P* | NRQ | *P* | NRQ | *P* | NRQ | *P* |
| **LGG CFS** | SC5314 | 0.476 ± 0.054 | **0.0030** | 0.718 ± 0.184 | 0.2546 | 1.202 ± 0.174 | 0.3590 | 0.594 ± 0.087 | **0.0271** | 0.655 ± 0.124 | 0.0615 | 0.943 ± 0.146 | 0.6223 |
|  | A14 | 0.573 ± 0.068 | **0.0104** | 0.675 ± 0.121 | 0.0697 | 0.996 ± 0.095 | 0.8976 | 0.729 ± 0.109 | 0.1076 | 0.694 ± 0.031 | **0.0012** | 1.213 ± 0.269 | 0.5191 |
|  | A69 | 0.843 ± 0.102 | 0.1887 | 0.849 ± 0.126 | 0.2958 | 0.789 ± 0.064 | **0.0379** | 0.796 ± 0.138 | 0.2115 | 0.706 ± 0.049 | **0.0066** | 1.295 ± 0.296 | 0.3889 |
|  | T18S | 0.629 ± 0.150 | 0.0798 | 0.469 ± 0.110 | **0.0406** | 0.786 ± 0.307 | 0.4373 | 0.571 ± 0.156 | 0.1206 | 1.113 ± 0.432 | 0.8952 | 0.313 ± 0.146 | 0.0981 |
|  | T38R | 1.648 ± 0.119 | **0.0027** | 0.742 ± 0.069 | **0.0308** | 1.216 ± 0.119 | 0.1453 | 1.080 ± 0.066 | 0.2885 | 1.710 ± 0.096 | **0.0007** | 0.831 ± 0.129 | 0.2787 |
| **LP8014 CFS** | SC5314 | 0.583 ± 0.047 | **0.0029** | 0.314 ± 0.080 | **0.0082** | 0.783 ± 0.165 | 0.2598 | 0.562 ± 0.142 | 0.0584 | 0.791 ± 0.186 | 0.3267 | 0.434 ± 0.070 | **0.0064** |
|  | A14 | 0.948 ± 0.155 | 0.6404 | 0.862 ± 0.084 | 0.1733 | 1.137 ± 0.029 | **0.0071** | 0.786 ± 0.019 | **0.0005** | 1.067 ± 0.108 | 0.6407 | 1.161 ± 0.170 | 0.4365 |
|  | A69 | 1.176 ± 0.190 | 0.4526 | 0.593 ± 0.134 | 0.0629 | 0.812 ± 0.086 | 0.1070 | 0.764 ± 0.148 | 0.2222 | 0.887 ± 0.082 | 0.2243 | 1.048 ± 0.242 | 0.9855 |
|  | T18S | 0.664 ± 0.157 | 0.1052 | 0.440 ± 0.100 | **0.0299** | 0.819 ± 0.262 | 0.4574 | 0.736 ± 0.151 | 0.2000 | 1.279 ± 0.365 | 0.6558 | 0.360 ± 0.170 | 0.0956 |
|  | T38R | 1.770 ± 0.096 | **0.0004** | 0.700 ± 0.036 | **0.0021** | 0.954 ± 0.048 | 0.3908 | 0.930 ± 0.079 | 0.4156 | 1.342 ± 0.099 | **0.0186** | 0.609 ± 0.095 | **0.0373** |

\*Expression levels are expressed as mean ± SEM of the normalized relative quantity (NRQ) obtained from triplicate experiments. NRQ of control group (MRS) is constantly 1.0 and not shown. †P-value was obtained by comparing to the control group using Student's t test. Bold values denote statistical significance at $p < 0.05$ .

## 4 Discussion

Lactobacillus species have been reported to exhibit an inhibitory effect on biofilm formation and filamentation of *C. albicans*, however few studies have described their effect on the non-albicans species. In the present study, we studied the antibiofilm effect of cell suspensions and CFSs of three *Lactobacillus* strains on *C. albicans, C. tropicalis*, and *C. parapsilosis*. LGG CFS shown the strongest anti-biofilm activity against *C. albicans* and *C. tropicalis* strains, followed by LP8014 CFS. The *C. parapsilosis* were more susceptible to LA4356 than LGG and LP8014. Previous studies have reported the anti-biofilm activity of *Lactobacillus* cell suspensions on *C. albicans*, however, we detected only a modest effect of cell suspension on *Candida* biofilm in our study(Vilela et al., 2015; Matsubara et al., 2016b; Ribeiro et al., 2017). It is noteworthy that different methods of preparing the cell suspensions were used. In our study, *Lactobacillus* cells were harvested and resuspended in fresh media, while cell suspensions used in other studies were directly diluted from a pre-incubated liquid culture to the desired concentration. Vilela *et al*. has tested the antibiofilm activity of *L. acidophilus* culture at different growth phases (4, 6, 18 and 24 hour) on *C. albicans* biofilms and found that the reduction of *C. albicans* CFU in biofilm compared to control was only statistically significant for the 24-hour *L. acidophilus* culture, suggesting that inhibitory activity of *Lactobacillus* is possibly associated with the growth stage(Vilela et al., 2015). In the present study, *C. parapsilosis* was found to be less susceptible to CFSs of LGG and LP8014. Similar *Candida* species-specific manner of the *Lactobacillus* inhibitory activity was reported in previous research. Tan *et al*. observed that biofilm formation of *C. parapsilosis* was less suppressed by the supernatant of *L. gasseri* and *L. rhamnosus* than *C. tropicalis* and *C. krusei*(Tan et al., 2018). Parolin *et al*. investigated the antifungal activities of CFS of *Lactobacillus* strains isolated from vaginal swabs on *C. albicans* and non-albicans species but found that no strains were effective against *C. krusei* and *C. parapsilosis*, even when being effective on other *Candida* species(Parolin et al., 2015).

Our study showed an inhibitory effect of *L. rhamnosus* (LGG) and *L. plantarum* (LP8014) CFSs on *Candida* biofilm. This finding corroborates the results of numerous studies using other *Lactobacillus* species and strains(Vilela et al., 2015; James et al., 2016; Matsubara et al., 2016b; Ribeiro et al., 2017; Matsuda et al., 2018; Tan et al., 2018). The indirect interaction between *Lactobacillus* cells and *Candida* biofilm suggests that soluble exometabolites secreted into the culture medium by *Lactobacillus* are possibly responsible for the inhibitory effect. Production of lactic acid and the resulted low pH (4.5) environment has been accounted for candicidal activity of *L. rhamnosus* and *Lactobacillus reuteri* (Köhler et al., 2012). However, in our biofilm assay, the lowest pH of medium measured was 6.0 and caused by the addition of LGG CFS. Moreover, acidified MRS broth showed little inhibitory effect on biofilm, and the inhibitory effect was not diminished in neutralized LGG CFS (p <0.05 when compared with MRS control, except for biomass of *C. tropicalis* T18S and T38R) and only reduced when compared with untreated LGG CFS in *C. albicans* A14 and A69. These results suggested that there are other factors contributed to the inhibitory effect.

*Lactobacillus* have been described to produce other exometabolites with antimicrobial and antibiofilm effects, such as small chain-fatty acids, H_2_O_2_, antimicrobial peptides, and biosurfactants(Strus et al., 2005; Gudiña et al., 2010; Nguyen et al., 2011; Rushdy and Gomaa, 2013; Ceresa et al., 2015). Sodium butyrate, a small chain fatty acid produced by *Lactobacillus* species, is a known histone deacetylase inhibitor(Garnaud et al., 2016). Nguyen *et al*. demonstrated that sodium butyrate can significantly inhibit biofilm formation of *C. albicans* and *C. parapsilosis*, by 80% and 65% respectively, at a concentration of 10 mM(Nguyen et al., 2011). Song and Lee investigated the antifungal activity of spent culture medium of *L. rhamnosus* and *L. casei* and found that the growth of *C. albicans* in both yeast- and hyphal-predominant conditions were significantly inhibited with 50% v/v spent culture medium(Song and Lee, 2017). However, proteinase K-treated spent culture medium of both species lost the inhibitory effect, suggesting that the effect was due to antifungal peptides. A biosurfactant produced by a *Lactobacillus brevis* strain CV8LAC was shown to inhibit *C. albicans* biofilm formation by coincubation and precoating the substrate whiles demonstrated no inhibition on the fungal growth in planktonic and sessile form (Fracchia and Allegrone, 2010; Ceresa et al., 2015). It is possible that these metabolites and similar compounds also contribute to the inhibitory effect of CFS demonstrated in our study, but further investigation is needed to identify the exact active compounds and their mechanisms.

Filaments are key structural component of mature biofilm of *C. albicans*(Chandra et al., 2001; Kuhn et al., 2002). Mutants of hyphal-related genes showed defects in forming hyphae and biofilm formation as well(Ramage et al., 2002; Nobile and Mitchell, 2005). Therefore, we investigated the effect of CFSs of LGG and LP8014 on the filamentation of the *C. albicans* and *C. tropicalis* strains. The filamentation of *C. albicans* and *C. tropicalis* cells was greatly reduced when co-incubated with the CFSs. These results are consistent with other studies that demonstrated inhibitory effects of *Lactobacillus* cell suspensions and CFSs on *C. albicans* in filamentation assay of similar designs (Vilela et al., 2015; Ribeiro et al., 2017; Matsuda et al., 2018; Rossoni et al., 2018b). Matsubara *et al*. observed a significantly reduction of hyphal elements in *C. albicans* biofilm treated with *L. rhamnosus* supernatant by confocal laser scanning microscopy(Matsubara et al., 2016b). In a monolayer model of oral epithelial cells, preincubating the epithelial cells with LGG suspension for 12 h before *C. albicans* infection was shown to significantly reduce hyphal length and invasion of *C. albicans* (Mailänder-Sánchez et al., 2017).

Environmental pH is known to induce morphological differentiation in *C. albicans*. Growth in yeast from is favored in acidic conditions, whereas neutral and alkaline conditions prompt hyphal growth(Davis, 2003). We found that the addition of LGG and LP8014 CFSs reduced the pH of the medium from 7.5 of the control group (MRS broth) to between 6 and 7. As Nadeem *et al*. has observed that germ tube formation in *C. albicans* at pH 6.4 was moderately lower than at pH 7.4 and further reduced at pH 5.4, we could not exclude the effect of the slightly decreased pH in the inhibition of filamentation observed in our study, but other metabolites of *Lactobacillus* are suggested to contribute(Nadeem et al., 2013). For example, in a recent study, 1-acetyl-β-carboline (1-ABC), a small molecule isolated from the culture supernatant of *Lactobacillus* species, was reported to block *C. albicans* filamentation in a concentration-dependent manner(MacAlpine et al., 2021). It was proposed that 1-ABC inhibits *C. albicans* Yak1 kinase, which is necessary for upregulation of hypha-induced genes(Goyard et al., 2008). Purified 1-ABC was also shown to inhibit filamentation of *C. dubliniensis* and *C. tropicalis*, in which Yak1 is conserved. The authors also demonstrated that acidic pH alone was insufficient to achieve the same degree of filamentation inhibition caused by *Lactobacillus* culture supernatant.

To elucidate the mechanisms of the antibiofilm activity of *Lactobacillus* CFS, we measured the expression of six biofilm-related genes, namely *ALS1, ALS3, BCR1, EFG1, TEC1*, and *UME6*, in the biofilm produced by *C. albicans* or *C. tropicalis* co-incubated with LGG and LP8014 CFSs. *ALS1* and *ALS3*, belonging to the agglutinin-like sequence (*ALS)* gene family, encode cell wall glycoproteins Als1 and Als3 respectively(Hoyer, 2001). Expression of *ALS1* is detectable in both yeast or hyphae form, whiles *ALS3* transcribed exclusively in germ tubes and hyphae(Green et al., 2005). In *C. tropicalis*, Galán-Ladero *et al*.(Galán-Ladero et al., 2019) reported that *ALS1*-, *ALS2*-, and *ALS3*-like genes were more upregulated in sessile cells, suggesting their roles in biofilm formation. *BCR1*, encoding a transcription factor Bcr1, is upregulated in *C. albicans* hyphae but not required for normal hyphal development(Nobile and Mitchell, 2005). Bcr1 is required for the full expression of several cell wall proteins, including Als1 and the hyphal-specific Als3, Hwp1, and Hyr1. *EFG1* encodes the transcription factor Efg1, which is essential for yeast-hyphal transition and biofilm formation of *C. albicans*(Stoldt et al., 1997; Ramage et al., 2002). Efg1 is activated by upstream cAMP pathway and regulates expression of hyphal-specific genes and downstream transcription factors such as Tec1 and Eed1(Sudbery, 2011). Mancera *et al*. has reported that *EFG1* ortholog in *C. tropicalis* has conserved function on filamentation and biofilm formation(Mancera et al., 2015). *TEC1* encodes transcription factor Tec1, deletion of which results in defect of hyphal formation and biofilm formation, and suppression of expression of secreted aspartyl protease (SAP) family proteins Sap4-6 in *C. albicans*(Schweizer et al., 2000; Nobile and Mitchell, 2005). In *C. tropicalis*, deletion mutants of *BCR1, BRG1, TEC1, EFG1*, or *NDT80* homologs produced significantly less biofilm biomass, suggesting the conserved roles of these genes in biofilm formation(Tseng et al., 2020). *UME6*, a common downstream target of regulators Efg1, Chp1 and Ras1, strictly or partially controls the expression of hyphae-specific genes in *C. albicans*(Zeidler et al., 2009). Constitutive high-level expression of *UME6* orthologs in *C. tropicalis* and *C. parapsilosis* also enhance filamentation and biofilm formation(Lackey et al., 2013).

To the best of our knowledge, this is the first study to examine the expression of biofilm-related genes in *C. tropicalis* biofilm under a biofilm-inhibitory condition. In our study, we observed a downregulation in *EFG1* and *TEC1* genes by CFS in the three *C. albicans* strains, which is in agreement with the findings of previous studies on *C. albicans* biofilm(James et al., 2016; Matsuda et al., 2018; Rossoni et al., 2018a). These studies, however, also reported significant downregulation of *ALS3* and *BCR1* genes in *Lactobacillus*-inhibited biofilm, which was not the case in our results. Interestingly, in our study, the expression of *ALS1, BCR1*, and *TEC1* genes were upregulated in *C. tropicalis* T38R by LGG CFS although LGG CFS was shown to suppress biofilm formation and filamentation of T38R, suggesting a strain-dependent gene response. The upregulation of *ALS1* in T38R could possibly be a compensatory upregulation, which has been demonstrated in deletion mutants of *ALS2* and *ALS4*, and of the *SAP* gene family in *C. albicans*, for the downregulation of *ALS3*(Zhao et al., 2005; Naglik et al., 2008; Liu and Filler, 2011). The expression of *UME6* in *C. albicans* SC5314, *C. tropicalis* T18S and T38R was downregulated by 0.434-, 0.360-(p=0.0956), and 0.609-fold respectively, suggesting that the mechanism of action of LP8014 CFS involves the repression of *UME6*. We selected few target genes from the biofilm regulatory network of *Candida*, which involves multiple signaling pathways and more than 50 genes(Finkel and Mitchell, 2011). Other well-known biofilm-related genes, such as *HWP1*, which encodes a cell wall adhesin required in biofilm formation, and *NRG1*, a negative regulator of filamentation, were not included(Braun et al., 2001; Nobile et al., 2006). To unravel the elusive mechanisms of the effect of *Lactobacillus* CFS, genomic-wide transcripting profiling by technologies such as RNA-seq is a better approach(Chong et al., 2018).

## 5 Conclusion

The present study evaluated the inhibitory effect of three lactobacilli strains, LGG, LP8014, and LA4356, on *C. albicans, C. tropicalis*, and *C. parapsilosis*. Significant inhibitory effects were demonstrated by the CFSs of LGG and LP8014 on filamentation and biofilm formation of both *C. albicans* and *C. tropicalis*. Hypha-related gene expressions in biofilm were also changed in response to the addition of CFSs. However, *C. parapsilosis* biofilms were not susceptible to the inhibitory effect of these two lactobacilli strains. The pathogen-specific inhibition suggests a common mechanism of action, presumably mediated by exometabolites of the lactobacilli, on *C. albicans* and *C. tropicalis*. This study demonstrated an alternative approach to control *Candida* biofilm. The active components and the potential clinical application of LGG and LP8014 CFSs remain to be investigated.

## Notes

### Competing Interest Statement

The authors have declared no competing interest.

