## Supplemental Table 1 for "Inhibitory effect of lactobacilli supernatants on biofilm and filamentation of *C. albicans, C. tropicalis*, and *C. parapsilosis*"

Table S1. Primers used in qPCR for *C. albicans.*

| **Species** | **Target** | **Primers** | **Sequence (5' to 3')** | **Product size (bp)** |
| --- | --- | --- | --- | --- |
| *C. albicans* | *ACT1* | AACT1 1F | F: TGGTGTTACTCACGTTGTTCCA | 100 |
|  |  | AACT1 1R | R: GGACAAATGGTTGGTCAAGTCTC |  |
|  | *ALS1* | AALS1 2F | F: ACCAGTATCATTCCATCATTTTCCC | 71 |
|  |  | AALS1 2R | R: TCAAATGTTGACAAATCGGAGGTT |  |
|  | *ALS3* | AALS3 2F | F: ATTCGATCCTAACCGCGACA | 119 |
|  |  | AALS3 2R | R: TTGGTGCAGTTTTGGTCAGGT |  |
|  | *BCR1* | ABCR1 3F | F: CCTCATCAAATGGGTGGTGGT | 110 |
|  |  | ABCR1 3R | R: GCCAATGGTTCGGGTCTTCT |  |
|  | *EFG1* | AEFG1 3F | F: GCACCAATCACCCCAAGTTC | 97 |
|  |  | AEFG1 3R | R: TTTGGCAACAGTGCTAGCTG |  |
|  | *TEC1* | ATEC1 2F | F: GATCCAGTGTCTTCTGTTGGC | 76 |
|  |  | ATEC1 2R | R: GGACACGTGAATAAGCACCA |  |
|  | *RIP1* | RIP1F | F: TGTCACGGTTCCCATTATGATATTT | 72 |
|  |  | RIP1AR | R: TGGAATTTCCAAGTTCAATGGA |  |
|  | *UME6* | AUME6 4F | F: GGTGTCTCTTCTGATGTTGGT | 106 |
|  |  | AUME6 1R | R: ACTTCCAGATCCTGTACCACT |  |
