## Supplemental Table 2 for "Inhibitory effect of lactobacilli supernatants on biofilm and filamentation of *C. albicans, C. tropicalis*, and *C. parapsilosis*"

Table S2. Primers used in qPCR for *C. tropicalis*.

| **Species** | **Target** | **Primers** | **Sequence (5' to 3')** | **Product size (bp)** |
| --- | --- | --- | --- | --- |
| *C. tropicalis* | *ACT1* | TACT1 3F | F: TACGCTGGTTTCTCCTTGCC | 116 |
|  |  | TACT1 3R | R: GCGGTGGTGGAGAAAGTGT |  |
|  | *ALS1* | TALS1 2F | F: TCTGTTGCCATTCCTGTCGAA | 70 |
|  |  | TALS1 2R | R: AAACCAACCCAAGCAAGATCG |  |
|  | *ALS3* | TALS3 2F | F: TGACTCCTAAGGAAGTATCGGGA | 102 |
|  |  | TALS3 2R | R: GCCAAGCTGGATTAGCAGGA |  |
|  | *BCR1* | TBCR1 3F | F: ATCCTTTACCTGCGTTGCGT | 146 |
|  |  | TBCR1 3R | R: GGACGAAGCTACTGACTCGG |  |
|  | *EFG1* | TEFG1 3F | F: TCCTGCAGCTCCACCTATACC | 99 |
|  |  | TEFG1 3R | R: TTGGCTGGGTTTAACGTGTCT |  |
|  | *TEC1* | TTEC1 3F | F: TTCAAGCAGTGCTCCAAGTTC | 103 |
|  |  | TTEC1 3R | R: AGGTGCAGAATGGAAACCAGT |  |
|  | *RIP1* | RIP1F | F: TGTCACGGTTCCCATTATGATATTT | 72 |
|  |  | RIP1TR | R: TGGGATTTCCAAGTTCAATGGA |  |
|  | *UME6* | TUME6 3F | F: GGTTCAGCTCCAACAGAGACA | 131 |
|  |  | TUME6 3R | R: CCTGGTTGGCTGTATTCTCCA |  |
